# AN INULIN-TYPE FRUCTAN ENRICHED EXCLUSIVE ENTERAL NUTRITION FORMULA SUPPRESSES COLITIS THROUGH GUT MICROBIOME MODULATION AND PROMOTING EXPANSION OF ANTI-INFLAMMATORY T CELL SUBSETS

**DOI:** 10.1101/2021.02.02.429403

**Authors:** Genelle R. Healey, Kevin Tsai, Daniel J. Lisko, Laura Cook, Bruce A. Vallance, Kevan Jacobson

## Abstract

**Background & Aims:** Exclusive enteral nutrition (EEN) is used to treat pediatric Crohn’s disease (CD), but therapeutic benefits are not long lasting. Due to reported lower efficacy EEN is not routinely used to treat pediatric ulcerative colitis (UC). Inulin-type fructans (IN) beneficially modulate the gut microbiome and promote expansion of anti-inflammatory immune cells. We hypothesized that enriching EEN with IN (EENIN) would enhance treatment efficacy. To test this, we examined the effects of EEN-IN on colitis development, the gut microbiome and CD4^+^ T cells using an adoptive T cell transfer model of colitis.

**Methods:** TCR-ß deficient mice were randomized to one of four groups: 1) Control, 2) Chow, 3) EEN and 4) EEN-IN, and naïve CD4^+^ T cells were adoptively transferred into groups 2-4, after which mice were monitored for 5-weeks prior to experimental endpoint.

**Results:** Mice fed EEN-IN showed greater colitis protection, with colonic shortening, goblet cell and crypt density loss reduced over that of EEN fed mice and reduced disease activity and immune cell infiltration compared to chow fed mice, and less crypt hyperplasia and higher survival compared to both groups. EENIN mice maintained colonic mucus layer thickness and had increased levels of Foxp3^+^IL-10^+^ and Rorγt^+^IL- 22^+^ and reduced levels of Tbet^+^IFNγ^+^ and Tbet^+^TNF^+^ CD4^+^ T cells. EEN-IN also lead to higher butyrate, *Bifidobacterium* spp. and *Bacteroides* spp. concentrations.

**Conclusion:** The EEN-IN group showed reduced colitis development as compared to the chow and EEN groups. This highlights the potential benefits of EEN-IN as a novel induction therapy for pediatric CD and UC patients.

**Synopsis:** We demonstrated that inulin-type fructan enriched exclusive enteral nutrition formula reduced colitis development likely due to butyrate-dependent pathways that helped preserve the mucus layer and promote an anti-inflammatory intestinal environment via expansion of regulatory T cells.

## Introduction

Inflammatory bowel disease (IBD) is characterized by chronic, relapsing inflammation of the gastrointestinal (GI) tract. The etiology of IBD is unclear, but is thought to involve a complex interaction between genetics, the gut microbiome and other environmental triggers which lead to an aberrant immune response^1^. The most common forms of IBD are Crohn’s disease (CD), which can affect any part of the GI tract but is primarily found in the terminal ileum and cecum; and ulcerative colitis (UC), which affects only the colon. This idiopathic, relapsing and remitting disease has no cure or universally efficacious treatment, therefore, novel therapeutic strategies to reduce the burden of disease are in high demand.

Exclusive enteral nutrition (EEN), a nutritionally complete liquid diet with no food intake, is considered a gold standard treatment for inducing disease remission in newly diagnosed pediatric CD patients^2,3^. It provides comparable benefits over standard medications (i.e. corticosteroids and 5-aminosalicylates agents) but additionally promotes mucosal healing and improved nutritional status, as well as bone health and linear growth^4^. The mechanism(s) by which EEN achieves these benefits is unknown, but changes in the gut microbiota^5^, inflammatory mediators^6,7^ and intestinal barrier function^8^ are likely implicated. EEN is not routinely used in active UC patients as it is less efficacious in this patient group^9^. A reason for the lower efficacy may be related to the lack of indigestible substrates (i.e. fiber or prebiotics) found in the EEN formula predominately used; therefore, by the time the EEN reaches the inflamed colon of UC patients, most of the beneficial nutrients have already been absorbed in the small bowel. Even among CD patients, the benefits of EEN are variable, as the treatment is often associated with a dysbiotic gut microbiota profile^10–12^ and after EEN is stopped the reduction in fecal calprotectin levels (marker of intestinal inflammation) is generally lost^13,14^. Furthermore, up to 85% of CD patients experience disease relapse within 1-2 years following discontinuation of EEN^15–17^. In addition, studies have shown that dietary re-introduction after EEN leads to a major rebound in the abundance of Proteobacteria; a bacterial phylum often associated with inflammation^14^. Moreover, a low fiber diet following EEN is associated with lower microbial diversity and increased risk of disease relapse^18^. We suspect the absence of prebiotics and fiber in EEN formula may be responsible for its suboptimal efficacy and outcomes in children with IBD. Hence, the addition of fiber or prebiotics to EEN formula might increase its efficacy, especially in more distal disease, and may prevent EEN associated dysbiosis.

Previous studies have demonstrated that prebiotics and fiber beneficially modulate the gut microbiome and reduce inflammation in IBD^19–21^. Inulin-type fructans (IN) appear to be particularly promising in enhancing clinical outcomes in adult IBD patients, suppressing inflammation as well as modulating the composition and function of the gut microbiome^22,23^. However, to date, no studies have investigated the therapeutic efficacy of an IN enriched EEN (EEN-IN) using animal models of colitis or in IBD patients. We hypothesize that the addition of IN to EEN will lead to beneficial modulation of the gut microbiome and promote the expansion of anti-inflammatory CD4^+^ T cells which will lead to suppression of colitis as compared to the chow and EEN groups. Thus, we investigated the potential anti-inflammatory and gut microbiome modulating properties of a novel EEN-IN formula, as compared to fiber-free EEN and chow, using a well-established adoptive T cell transfer model of murine colitis.

## Results

### Mice fed EEN-IN were protected from developing colitis

As measured by DAS, there was significantly more colitis development in the chow group as compared to the control and EEN-IN groups, due to more chow fed mice experiencing weight loss and diarrhea. Colitis development was also lower in the EEN-IN group compared to the EEN group, however, this difference didn’t reach significance (Figure 1A). The EEN fed mice had a higher body weight change at the end of the experiment when compared to all other groups (Figure 1B). This is likely explained by the higher calorie consumption observed in the EEN group (Table 1). Moreover, all control and EEN-IN mice survived until the end of the experiment, however, survival rates were only 92% and 77% in the EEN and chow groups; respectively, as several mice in these groups had to be euthanized early as they reached humane endpoint due to the development of severe colitis (Figure 1C). While EEN and chow fed mice had increased spleen weights (indicating more severe colitis) compared to the control group, strikingly, spleen weights in the EEN-IN group were comparable to those of the control group. The EEN-IN group also had significantly lower spleen weights as compared to the chow group. Lower spleen weights in the EEN-IN fed mice suggest reduced systemic inflammation particularly in comparison to the chow fed mice (Figure 1D). EEN-IN fed mice also experienced significantly less colonic shortening compared to EEN fed mice, with colon lengths similar to control mice (Figure 1E).

**Figure 1.**
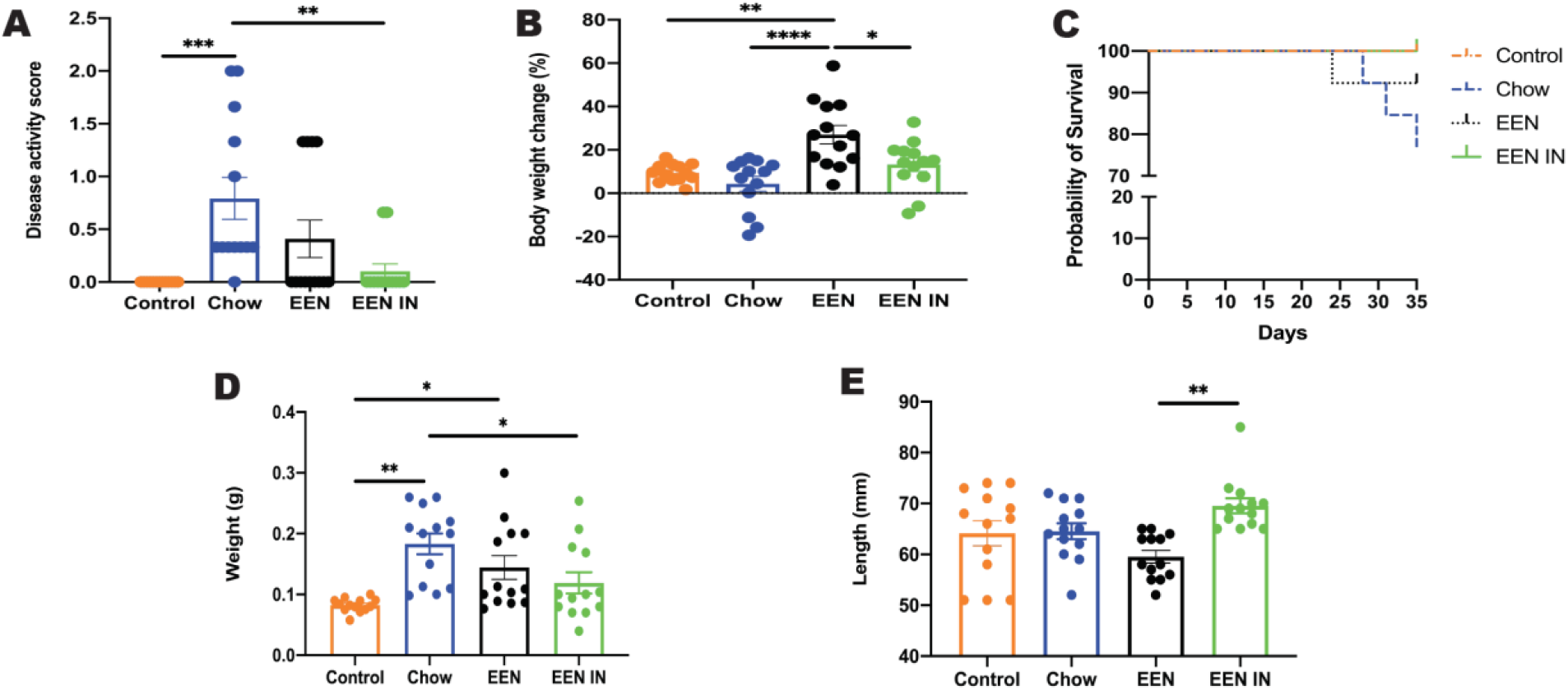
Mean (± SEM) (**A**) disease activity score (DAS), (**B**) body weight change, (**C**) survival rates, (**D**) spleen weight and (**E**) colon length between the four intervention groups (n = 13 mice/group; each data point represents one mouse). * p < 0.05, ** p < 0.01, *** p < 0.001, **** p < 0.0001.

**Table 1:** Nutritional composition of the experimental diets as well as the average intake of each diet per group.

|  | Control/Chow | EEN | EEN-IN |
| --- | --- | --- | --- |
| **Diet type** | Teklad TD.97184 (AIN-93G) | Ensure® Plus | Ensure® Plus and Orafiti® synergy 1 |
| **Energy (kcal/g or kcal/mL)** | 3.8 | 1.51 | 1.57 |
| **Protein (%)** | 18.8 | 15.1 | 14.6 |
| **Fat (%)** | 17.2 | 28 | 27.7 |
| **Carbohydrate (%)** | 63.9 | 56.9 | 57.9 |
| **Fibre (%)** | 5 | 0 | 3 |
| **Fibre sources** | Cellulose – poorly fermented | None | 50:50 inulin and FOS |
| **Average amount consumed (g or mL)** | 2.78 – Control<br>2.49 – Chow | 8.58 | 7.8 |
| **Average calories consumed (kcal/day)** | 10.56 – Control<br>9.46 – Chow | 12.96 | 12.24 |

EEN – standard exclusive enteral nutrition, EEN-IN – inulin-type fructan enriched exclusive enteral nutrition, FOS – fructo-oligosaccharide.

### Minimal histopathological changes and loss of mucus layer thickness observed in EEN-IN fed mice

H&E stained distal colonic sections were scored by two independent observers, to determine if differences in total and individual criteria scores were evident. Total histology scores were significantly higher in the chow and EEN as compared to the control group, whereas the EEN-IN group was not significantly different from the control group (Figure 2A). The chow and EEN groups were characterized by increased inflammatory cell infiltration, goblet cell and crypt density loss, crypt hyperplasia and submucosal inflammation compared to the control group. In contrast, only inflammatory cell infiltration was significantly elevated in the EEN-IN group as compared to the control group; no other histopathological evidence of colitis was observed (Figure 2A and B).

**Figure 2.**
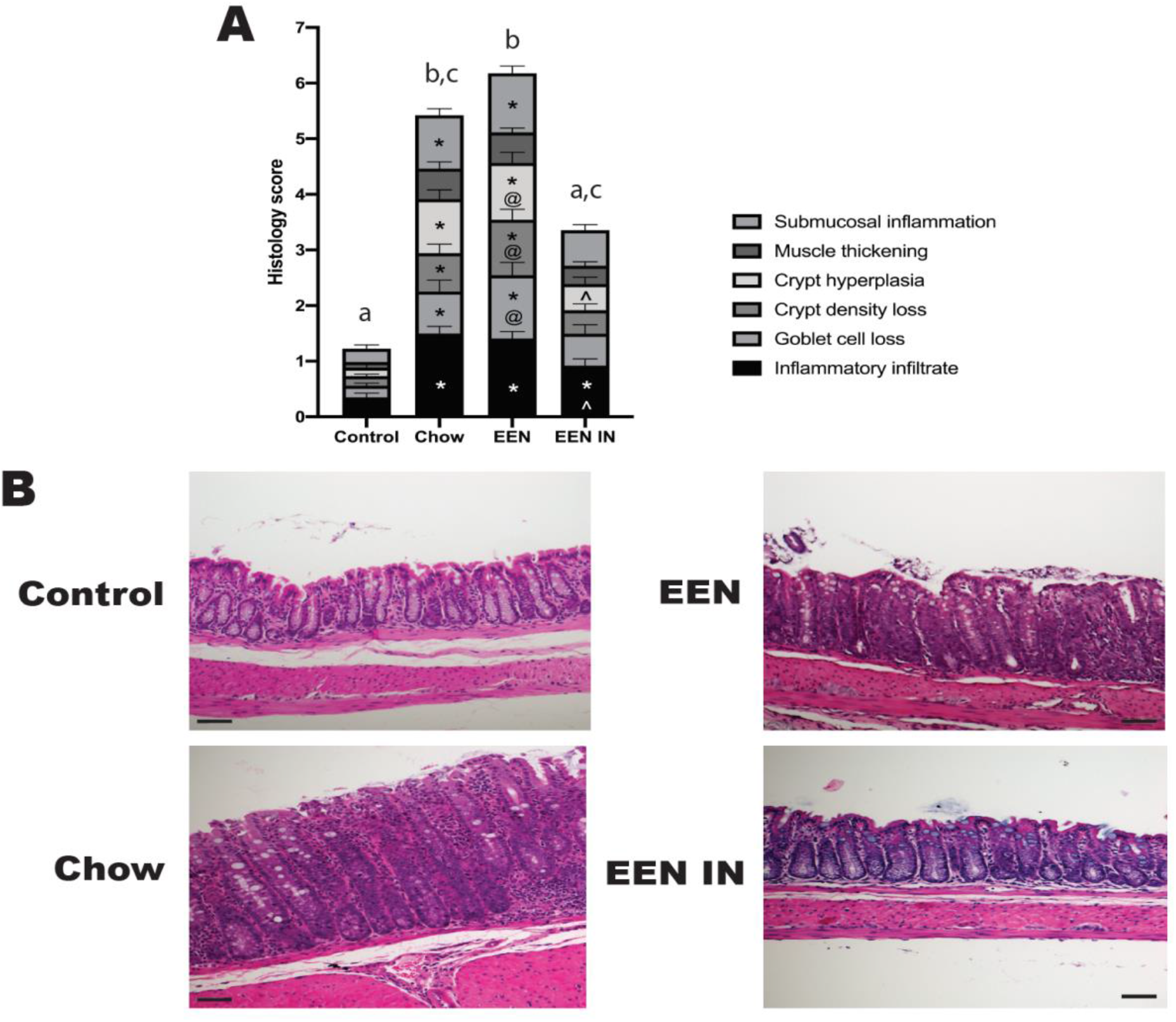
(**A**) A stacked bar graph showing the mean (± SEM) total histology scores as well as individual histology criteria score differences between the four intervention groups (n = 13 mice/group). Significant differences (p < 0.05) in total histology scores between groups are depicted by the different letters. Significant differences in individual histology criteria scores are depicted as follows: * control differs from chow, EEN or EEN-IN (p < 0.05), ^ chow differs from EEN-IN (p < 0.05), @ EEN differs from EEN-IN (p < 0.05). (**B**) Representative images of H&E stained distal colonic tissue from the control, chow, EEN and EENIN groups (x20; scale bar is 50 μm). Control – normal histopathology, chow and EEN – inflammatory infiltration, hyperplasia, submucosal inflammation, crypt density and goblet cell loss, EEN-IN – mild inflammatory infiltration.

Mid colonic sections were stained with alcian blue/PAS to investigate the effect each intervention had on goblet cell numbers and the mucus layer. Goblet cell numbers were significantly lower in the chow group as compared to the EEN-IN group (Figure 3A). While colitis in the chow and EEN groups both led to significantly thinner mucus layers, as compared to the control group, there was no significant reduction in mucus layer thickness in the EEN-IN group (Figure 3B). Therefore, an IN enriched EEN formula helped preserve goblet cell numbers and minimized the deterioration in the mucus layer compared to chow and EEN fed mice. Additionally, it appears that acidic mucins were more prevalent in the EEN and EEN-IN groups (goblet cells stained blue), whereas the control and chow groups appear to have both acidic and neutral mucins (goblet cells stained a mix of magenta and purple) (Figure 3C).

**Figure 3.**
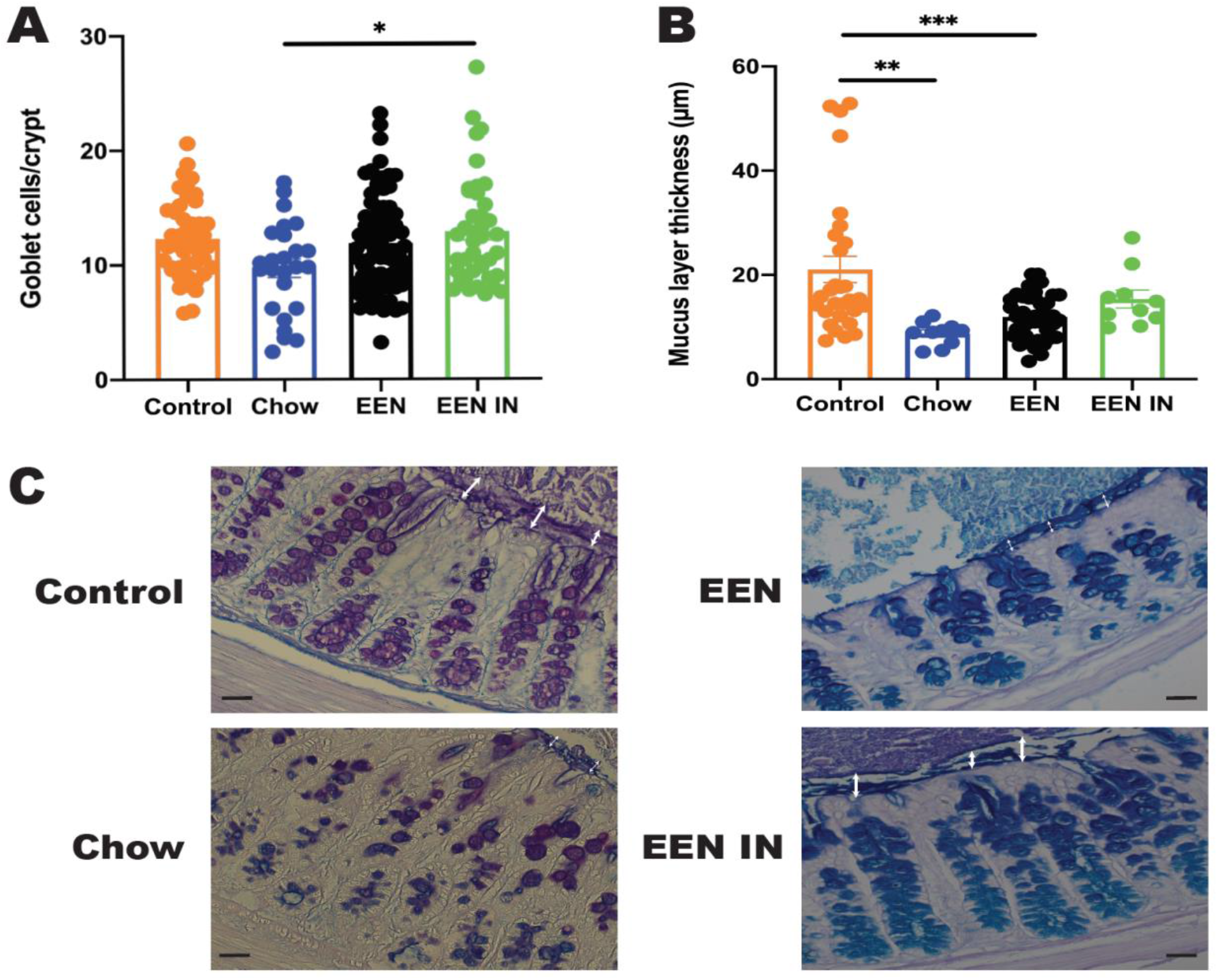
(**A**) Mean (± SEM) goblet cell numbers per crypt and (**B**) mucus layer thickness differences between the four intervention groups (n = 5 to 9 mice/group; each data point represents an individual measurement). **(C)** Representative images of alcian blue/PAS stained mid colonic sections from each intervention group (x40; scale bar is 40 μm). The white arrow depicts the mucus layer thickness for each group. Acidic mucins stained blue, neutral mucins stained magenta and a mix of acidic and neutral mucins stained blue/purple within goblet cells. * p < 0.05, ** p < 0.01, *** p < 0.001.

### EEN-IN results in reduced pro-inflammatory and expanded anti-inflammatory CD4^+^ T cells subsets

Next, we investigated whether there were any differences in the presence of pro-inflammatory and antiinflammatory CD4^+^ T cell subsets in the spleen and MLN between the chow, EEN and EEN-IN groups. As the control mice were αβ T cell deficient due to the absence of function of TCR-ß loci and were not adoptively transferred with naïve CD4^+^ T cells, CD4^+^ T cell numbers were very low compared to the other groups (0.25% versus 5.58%), therefore, these mice were not included in the CD4^+^ T cell subset analysis.

Chow fed mice were shown to have higher frequencies of the pro-inflammatory (Tbet^+^IFNγ^+^ and Tbet^+^TNF^+^) CD4^+^ T cell subsets in the spleen and MLN as compared to the EEN and EEN-IN groups (Figure 4B to E). While there were no significant differences in pro-inflammatory Rorγt^+^IL-17A^+^ CD4^+^ T cells in the spleen and MLN between groups (Figure 4F and G), there was a significant increase in frequency of antiinflammatory Foxp3^+^IL-10^+^ and Rorγt^+^IL-22^+^ CD4^+^ T cells in the spleen and MLN of mice fed EEN-IN compared to mice fed a chow diet (Figure 4H to K). Additionally, the frequency of splenic Rorγt^+^IL-22^+^ CD4^+^ T cells were also significantly increased in the EEN-IN fed mice compared to EEN fed mice (Figure 4H). Taken together, these data suggest that consumption of EEN-IN increased the frequency of anti-colitic CD4+ T cell subsets conferring protection against overt immune activation and colitis development.

**Figure 4.**
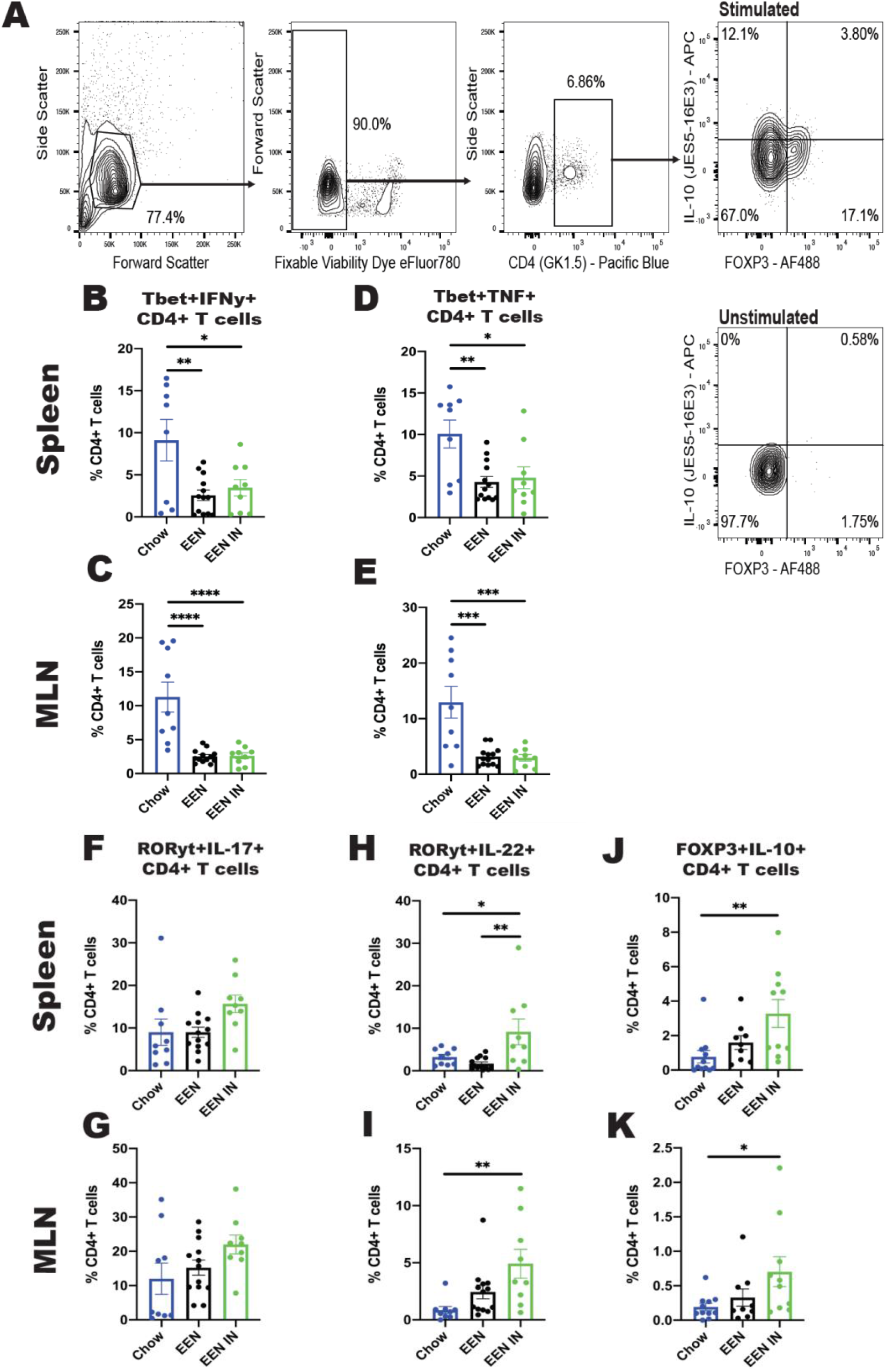
(**A**) Representative gating strategy for the enumeration of Foxp3^+^IL-10^+^ CD4^+^ T cells. Proportions of Tbet^+^IFNγ^+^ ([**B**] spleen and [**C**] MLN), Tbet^+^TNF^+^ ([**D**] spleen and [**E**] MLN), Rorγt^+^IL-17A^+^ ([**F**] spleen and [**G**] MLN), Rorγt^+^IL-22^+^ ([**H**] spleen and [**I**] MLN) and Foxp3^+^IL-10^+^ ([**J**] spleen and [**K**] MLN) CD4^+^ T cells in mice from the chow, EEN and EEN-IN groups (n = 8 to 13 mice per group; each data point represents one mouse). * p < 0.05, ** p < 0.01, *** p < 0.001, **** p < 0.0001.

### Increased abundance of beneficial microbes and concentrations of butyrate seen in EEN-IN fed mice

Several gut microbes known to be reduced in individuals with IBD, such as *Bifidobacterium* and *F. prausnitzii*^29^, were quantified to determine what impact the EEN-IN formula had on modulating these important microbial taxa. We observed that both EEN interventions (EEN and EEN-IN) led to a significant increase in total bacteria when compared to the control group (Figure 5A). The EEN-IN fed mice also experienced a significant bloom in *Bifidobacterium* (Figure 5B) and *Bacteroides* spp. (Figure 5C) when compared to the three other intervention groups. Lastly, *F. prausnitzii* concentrations increased significantly in the EEN fed mice (Figure 5D).

**Figure 5.**
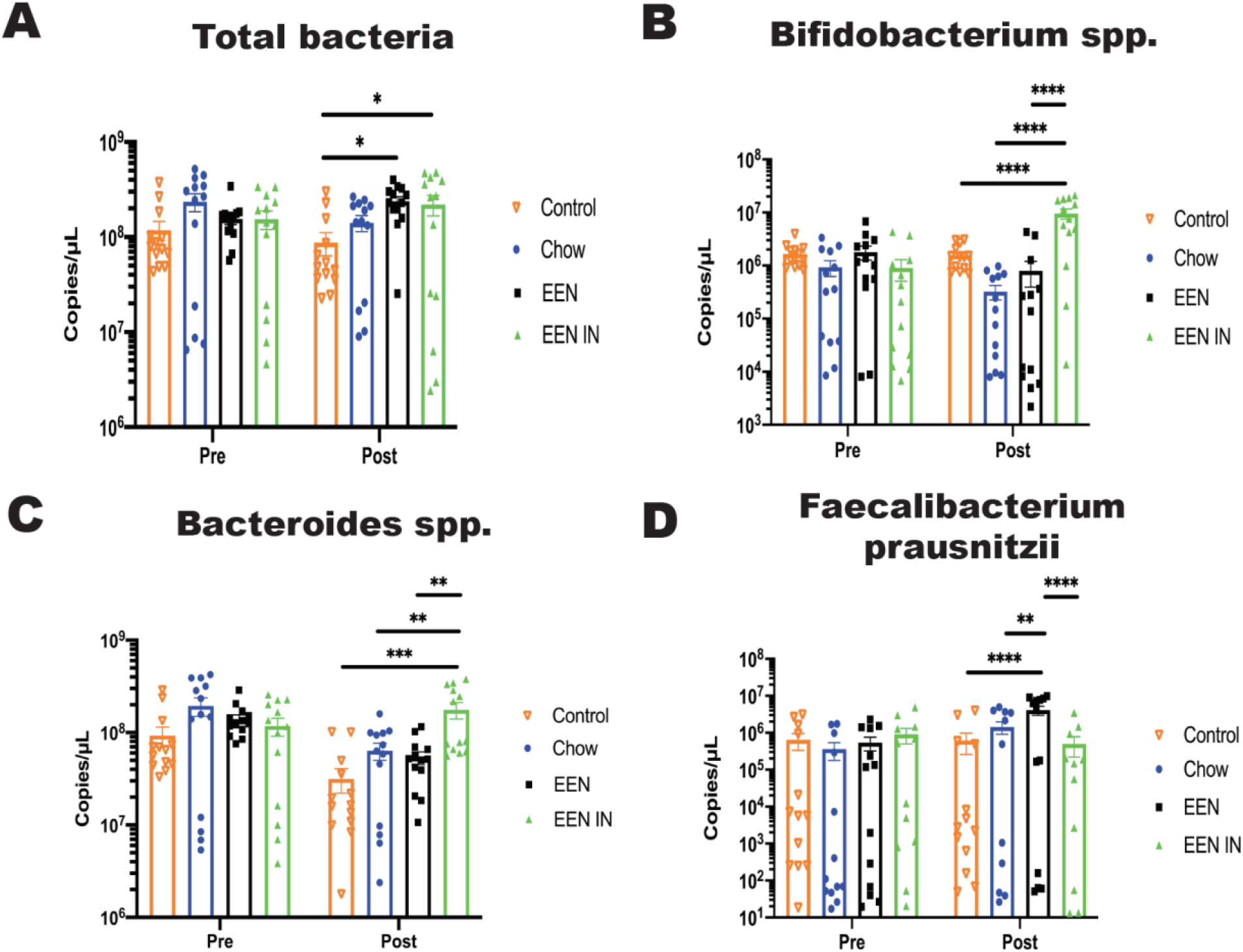
Mean (± SEM) (**A**) total bacteria, (**B**) *Bifidobacterium* spp., (**C**) *Bacteroides* spp. and (**D**) *Faecalibacterium prausnitzii* concentration (copies/μL) differences between the four intervention groups pre- and post-intervention (n = 13 mice/group; each data point represents one mouse). * p < 0.05, ** p < 0.01, *** p < 0.001, **** p < 0.0001.

Concentrations of certain microbial metabolites such as SCFAs, particularly butyrate, have also been shown to be in reduced in IBD patients. Therefore, we investigated whether the EEN-IN formula could enhance butyrate production. Indeed, cecal butyrate (Figure 6A) production was significantly higher in the EEN-IN group as compared to the chow and EEN groups. In contrast, neither cecal acetate (Figure 6B) nor propionate (Figure 6C) concentrations differed between any of the intervention groups. The branch-chain fatty acids (BCFA), isobutyrate (Figure 6D) and isovalerate (Figure 6E), were, however, found in significantly higher concentrations in the EEN group, but in significantly lower concentrations in the EEN-IN group, respectively. These data suggest that the addition of IN to EEN can lead to beneficial shifts in key microbial taxa, as well as a reduction in the concentrations of BCFAs and enhanced production of beneficial metabolites, such as butyrate.

**Figure 6.**
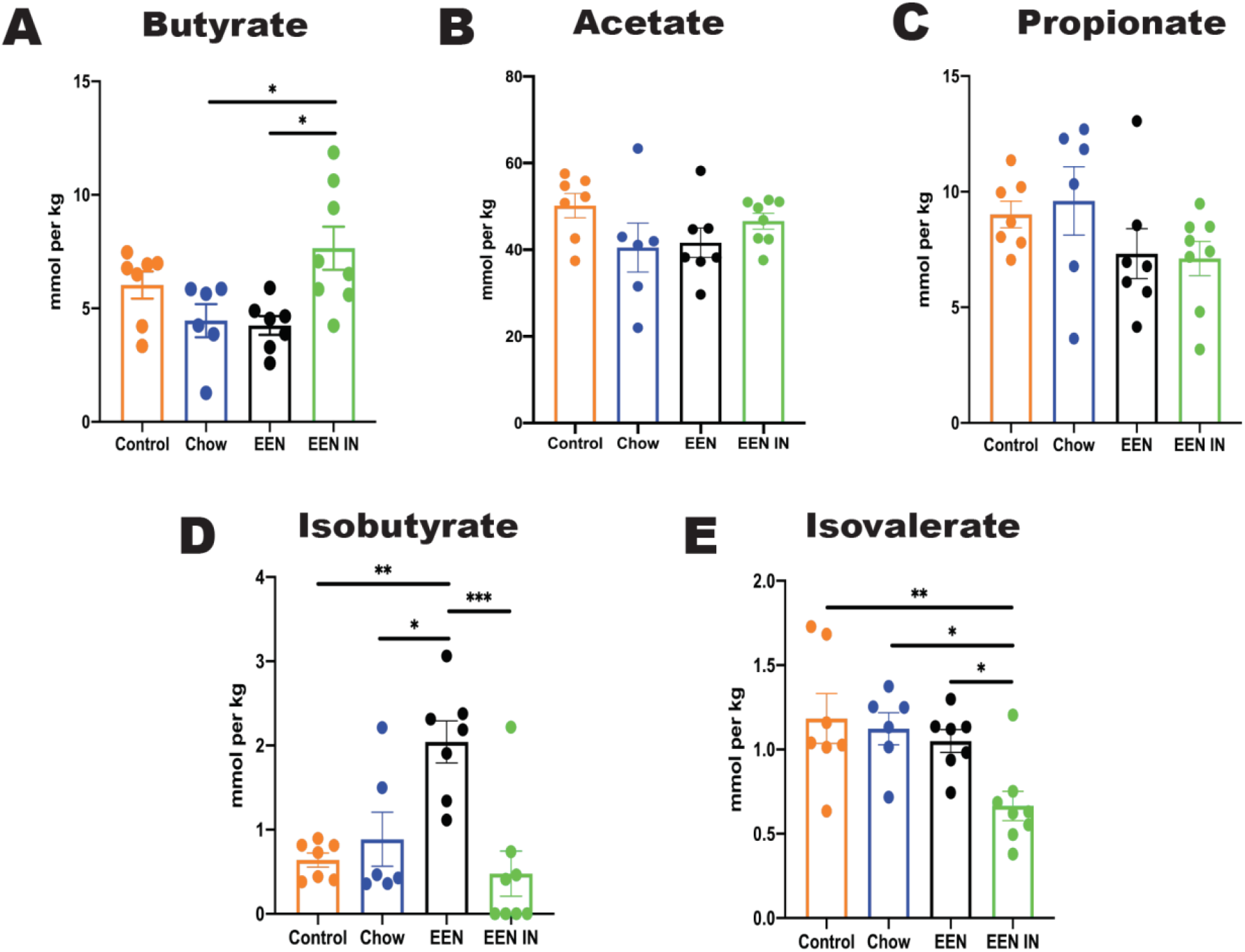
Mean (± SEM) cecal (**A**) butyrate, (**B**) acetate, (**C**) propionate (**D**) isobutyrate and (**E**) isovalerate concentration (mmoL/kg) differences between the four intervention groups (n = 6 to 8 mice/group; each data point represents one mouse). * p < 0.05, ** p < 0.01, *** p < 0.001.

## Discussion

In this study, we demonstrated that EEN-IN suppressed colitis development, with a concomitant expansion of anti-inflammatory CD4^+^ T cells and beneficial modulation of the gut microbiome. Specifically, consumption of an EEN-IN diet led to a 100% survival rate, as well as lower spleen weights, no colonic shortening and minimal histopathological changes in adoptive T cell transferred mice. The EEN-IN group also experienced less deterioration in mucus layer thickness along with higher colonic goblet cell counts. The gut microbiome of the EEN-IN fed mice was characterized by higher concentrations of *Bifidobacterium* and *Bacteroides* spp. as well as higher butyrate production. Lastly, lower proportions of pro-inflammatory CD4^+^ T cells and higher proportions of anti-inflammatory CD4^+^ T cells were observed in the spleen and MLN of EEN-IN fed mice.

Studies have shown that consumption of IN enhances health outcomes associated with several diseases^30,31^. One key property of IN is its ability to lead to a bloom in bacterial taxa such as *Bifidobacterium* and *Lactobacillus^32^* which, due to their ability to cross feed with butyrate-producing microbes, means IN also stimulates increased production of butyrate^33^. In fact, several of the health benefits associated with IN are thought to be related to its butyrogenic properties. Butyrate is a microbial metabolite that protects against several diseases such as colorectal cancer^34^, diabetes^35^, obesity^35^ and IBD^36^ via enhanced secretion of satiety hormones^37^, maintenance of intestinal barrier function^38^, improved insulin sensitivity^35^ and regulation of immune responses^39^.

One immune cell subset known to expand in the presence of high concentrations of butyrate are regulatory T cells (Tregs)^39^. Tregs are a sub population of CD4^+^ T cells essential for the maintenance of immune tolerance towards self-antigens and non-self-antigens^40^. Although most studies on Tregs focuses on their induction of xenograft tolerance, emerging evidence suggests that Tregs also induce tolerance towards resident microbiota as well as food antigens^41,42^. Despite these advancements, it is unclear whether Tregs can be induced by the microbiota and/or their metabolites through the administration of fermentable fibers such as IN. In this study we demonstrated that consumption of IN enriched EEN led to an expansion of Tregs, specifically Foxp3^+^IL-10^+^ CD4^+^ T cells, which suggests that fermentable fibers can induce Treg differentiation likely via a pathway dependent on IN stimulated butyrate production by resident gut microbes. Additionally, a key anti-inflammatory cytokine Tregs produce, IL-10, plays an important role in GI homeostasis with higher production of IL-10 leading to suppression of inflammation^24^. Several mechanisms have been proposed that link butyrate to the expansion of Tregs. Firstly, butyrate acts as a ligand upregulating peroxisome proliferator-activated receptor gamma (PPARγ) within intestinal epithelial cells which subsequently promotes Treg differentiation and survival by regulating the key Treg transcription factor, Foxp3^43^. The upregulation of PPARγ can also inhibit nuclear-factor-kappa B (NF-κB), a critical pro-inflammatory transcription factor, leading to a reduction in pro-inflammatory effector T cells^43,44^. Butyrate also binds with high affinity to G-protein coupled receptors (GPR) – GPR41, GPR43 and GRP109a. Butyrate bound GPRs promote antigen presenting cells (i.e. dendritic cells) to differentiate towards a more tolerogenic phenotype, which subsequently induces Treg expansion and increases IL-10 production^45^. Lastly, Foxp3 is also upregulated via butyrate-dependent histone deacetylase (HDAC) inhibition which potentiates the differentiation of naïve T cells to Tregs^39,46^.

Increased proportions of Rorγt^+^IL-22^+^ CD4^+^ T cells were also observed in mice fed EEN-IN. Similar to Tregs, the expansion of IL-22 producing CD4^+^ T cells is thought to be regulated by microbiota-derived butyrate via GPR41 binding and inhibition of HDAC. Enhanced IL-22 production has also been shown to suppress hyperactive immune responses in the intestine^47,48^. Microbiota-derived butyrate also has the capacity to regulate mucus production via an IL-22-dependent mechanism^49^. This is important as an intact mucus layer provides a barrier between the underlying intestinal epithelium and the microbes and food antigens within the lumen. Deterioration or weakening of the mucus layer can lead to encroachment of luminal microbes, leading to a pro-inflammatory response by intestinal epithelial cells and underlying immune cells. Additionally, butyrate, independent of IL-22, is able to enhance mucosal barrier integrity directly by stimulating mucin production by goblet cells^50^. Therefore, we propose that EEN-IN fed mice were protected from developing colitis via an IN-driven, butyrate-dependent expansion of antiinflammatory and potentially mucin stimulating T cell subsets (Foxp3^+^IL-10^+^ and Rorγt^+^IL-22^+^ CD4^+^ T cells) (Figure 7).

**Figure 7.**
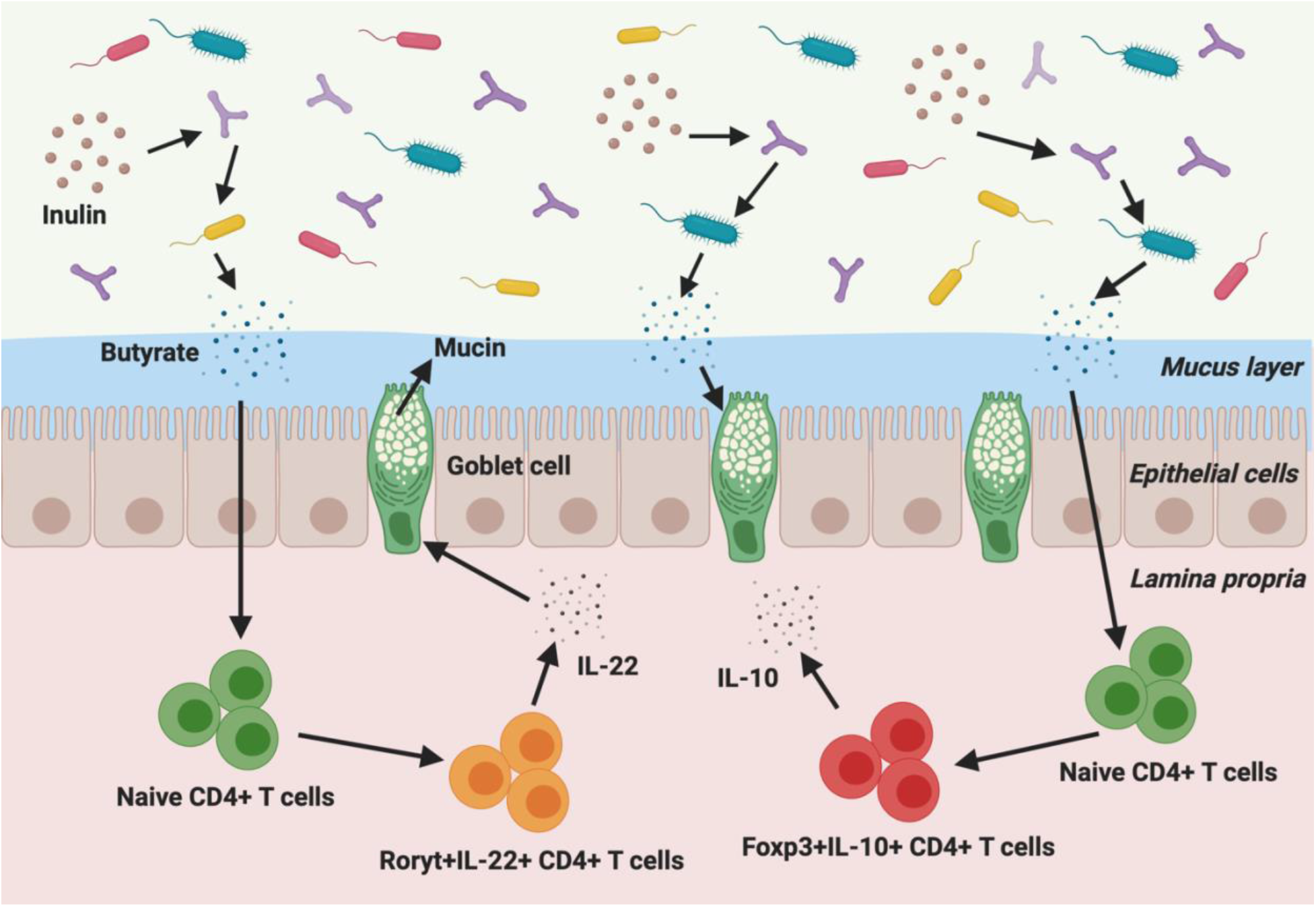
Schematic depicting the potential butyrate-dependent mechanistic pathways by which EEN-IN consumption can lead to reduced colitis development. IN provides the commensal microbes with a fermentable substrate, that via cross feeding, stimulates butyrate production. Butyrate promotes differentiation of naïve CD4^+^ T cells into anti-inflammatory CD4^+^ T cell subsets (i.e. Foxp3^+^IL-10^+^ and Rorγt^+^IL-22^+^ CD4^+^ T cells) which produce cytokines (i.e. IL-10 and IL-22) that reduce inflammation and lead to increased mucus production. Butyrate also has the ability to directly increase mucus production to maintain mucus layer thickness and enhance intestinal epithelial barrier function. Created with Biorender.com.

In contrast to EEN-IN fed mice, EEN and chow fed mice experienced colitis ranging from mild/moderate to severe. The EEN and chow groups both lacked fermentable fiber substrates which may at least partially explain the suboptimal outcomes observed in these groups as compared to the EEN-IN group. Previous research has shown that either chronic or intermittent reductions in dietary fiber causes the gut microbiota to utilize endogenous fermentable substrates, such as mucin glycoproteins within the intestinal mucus layer, as an energy source. A fiber-devoid diet subsequently leads to a shift in the gut microbiota from fiber-utilizing to mucus-eroding taxa. This leads to mucus layer deterioration and intestinal barrier dysfunction which promotes intestinal inflammation^51,52^. Dietary fiber deprivation also leads to a reduction in butyrate production^53^, impairing the beneficial immunomodulatory properties associated with this microbial metabolite (as outlined above). Therefore, the lack of available fermentable fiber in the EEN and chow diets likely promoted colitis development via deterioration of the mucus layer, alterations in gut microbiota composition, reduced butyrate production and an imbalance in pro- vs antiinflammatory CD4^+^ T cell subsets.

EEN is considered one of the gold standard therapies used to treat newly diagnosed pediatric CD patients, therefore we expected less severe colitis to develop in mice within the EEN group. However, the adoptive T cell transfer model is more reflective of human UC (colonic involvement only) rather than CD, therefore, these results may align more closely with the lower EEN efficacy observed in UC patients. Interestingly, we detected a bloom in *F. prausnitzii* within the EEN group, which was not observed in the EEN-IN or chow groups. IBD patients generally have reduced levels of *F. prausnitzii^29^,* with earlier studies suggesting that *F. prausnitzii* acts as an anti-inflammatory commensal, however, others have provided evidence that may discredit this claim. Specifically, it has been demonstrated that a reduction, rather than an increase, in *F. prausnitzii* abundance after EEN is correlated with improved clinical outcomes^54^. Therefore, it is possible that the increase in *F. prausnitzii* in the EEN group could either be a cause or consequence of colitis development. However, it’s unlikely that alterations in *F. prausnitzii* alone led to the colitic responses seen in the EEN group as *F. prausnitzii* concentrations were not significantly increased in the chow group. Both the chow and EEN groups did, however, contain similar levels of the SCFAs analyzed. As mentioned previously, the chow and EEN groups had lower butyrate concentrations as compared to the EEN-IN group. Additionally, the chow group had higher levels of the BCFA isovalerate and the EEN group had higher levels of both isovalerate and isobutyrate. BCFA, such as isovalerate and isobutyrate, are indicative of proteolytic fermentation^55^. Due to the lack of fiber in these diets it appears that the gut microbiota resorted to fermenting available amino acids leading to BCFA production. Little is known about the physiological role BCFAs play, but some studies suggest that increased protein fermentation and subsequent BCFA production may have detrimental effects on health^55,56^. Taken together, it appears that a lack of fiber in the chow and EEN diets led to thinning of the protective mucus layer and a shift towards a more dysbiotic gut microbiome profile which resulted in the proliferation of pro-inflammatory effectors CD4^+^ T cells and subsequent development of colitis.

## Conclusion

We can conclude that within an adoptive T cell transfer model of colitis, IN enriched EEN led to several favorable outcomes. EEN-IN was superior to EEN and chow as it suppressed colitis development, prevented loss in mucus layer thickness, increased butyrate production, promoted increased concentrations of *Bifidobacterium* and *Bacteroides* spp. and led to an expansion of anti-inflammatory T cell subsets, including IL-10 producing Tregs. We postulate that the enhanced outcomes observed in EENIN mice occurred via butyrate-dependent pathways that helped preserve the mucus layer and promoted an anti-inflammatory intestinal environment.

This study is the first animal or human study to investigate whether the additional of IN to EEN leads to enhanced outcomes compared to fiber-free EEN, the predominant formula used as an induction therapy in pediatric CD. Novel alternative or complementary pediatric IBD therapies with minimal side effects are currently in high demand, particularly for pediatric UC patients, where to date nutritional interventions have provided minimal benefit in helping induce remission. These data provide pre-clinical evidence to support undertaking human studies to investigate whether IN enriched EEN is a superior alternative option to fiber-free EEN for the treatment of both UC and CD patients.

## Materials and Methods

### Adoptive T cell transfer model of colitis

TCR-β deficient (-/-) C57BL/6 mice were bred at BC Children’s Hospital Research Institute animal facility under specific pathogen free conditions and were used at 8-14 weeks of age. A mix of both male and female mice were used, with a ratio of males to females of approximately 3:1 per group. We undertook six independent experiments using a total of 52 mice, with 13 mice randomly assigned to each of the four intervention groups: 1) Control group received a phosphate buffered saline (PBS) intraperitoneal (IP) injection and standard chow diet (negative control), 2) Chow group received a naïve T cell IP injection and standard chow diet (positive control), 3) EEN group received a naïve T cell IP injection and enteral formula without fiber and 4) EEN-IN group received a naïve T cell IP injection and enteral formula with fiber. Animal care, maintenance and experimental procedures were approved by the University of British Columbia Animal Care Committee (permit number: A19-0254) and performed in accordance with the Canadian Council on Animal Care (CCAC).

### Naïve T cell sorting and adoptive T cell transfer

Spleens and lymph nodes from female 8-12-week-old Thy1.1^+/+^ Foxp3-eGFP mice^24^ were processed into single cell suspensions by mashing the organs using a 40 μm nylon cell strainer. CD4^+^ T cells were isolated from the cell suspension by magnetic bead selection (Stem Cell), then stained for viability (eFluor780, Thermofisher) followed by surface staining using fluorescence conjugated anti-mouse CD4 (GK1.5, Thermofisher) and CD44 (IM7, Thermofisher). Naïve CD4^+^ T cells, defined as CD4^+^, CD25^neg^, CD44^low^ and Foxp3^neg/low^, were sorted by fluorescence activated cell sorting to a purity of 95%. The resulting naïve CD4^+^ T cells were washed twice with PBS, enumerated using a hemocytometer and resuspended in PBS to a concentration of 2.5 million cells/mL.

200 μL aliquot of naïve CD4^+^ T cells (0.5 million cells) or PBS (control group only) were intraperitoneally injected into the recipient TCR-β^-/-^ mice. This adoptive T cell transfer leads to a gut microbiota dependent expansion of naïve CD4^+^ T cells into effector CD4^+^ T cells which leads to significant colitis by 4-8 weeks post-injection^25^. Certain environmental factors (i.e. diet) and/or changes in the gut microbiome have the potential to limit colitis development by promoting the expansion of anti-inflammatory CD4^+^ T cell subsets and suppressing pro-inflammatory CD4^+^ T cell subsets.

### Diets

Mice were commenced on their respective diets immediately after naïve CD4^+^ T cell adoptive transfer. The control and chow groups were provided with *ad libitum* access to water (H_2_O) and to Teklad TD.97184 diet (Envigo, Madison, Wisconsin, USA, www.envigo.com); a standard irradiated chow with 5% cellulose (poorly fermented fiber source). The EEN mice were fed *ad libitum* fiber-free enteral formula (Ensure^®^ Plus, Abbott Laboratories, Chicago, Illinois, USA) in their H_2_O bottles. The EEN-IN mice were fed *ad libitum* enteral formula (Ensure^®^ Plus, Abbott Laboratories, Chicago, Illinois, USA) enriched with 3% IN prebiotic (Orafti^®^ Synergy 1, Beneo, Mannheim, Germany) in their H_2_O bottles (Table 1). The mice continued on their respective diets until the end of the experiment, at 5 weeks post adoptive T cell transfer.

### Body weight and food intake

Body weight and food intake were monitored daily. Change in body weight was reported as a percentage drop or gain in body weight from the original weight as measured on the day of adoptive T cell transfer.

### Disease activity scores

Disease activity was monitored in mice twice weekly until at least one mouse lost > 10% body weight and then disease activity was monitored daily. The disease activity scores (DAS) were determined using a previously reported scoring system^26^, based on percentage loss of body weight, stool consistency and signs of intestinal bleeding.

### Survival

If mice reached their humane endpoint prior to the end of the experiment, which was 5 weeks post adoptive T cell transfer, they were euthanized. Humane endpoint was reached if mice lost > 20% of their original body weight, had severe rectal excoriation due to diarrhea and/or had a pronounced decline in activity with a hunched appearance.

### Spleen weight and colon length

At the end of the experiment, mice were anesthetized with isoflurane and then euthanized via carbon dioxide asphyxiation and cervical dislocation. Spleens were removed, weighed and then stored on ice in PBS with 2% FBS and 2mM EDTA prior to flow cytometry analysis (as described in section 2.4). Colons were also removed, and their lengths recorded. Each colon was then cut, and a longitudinal section of the distal colon was taken for H&E staining (as described in section 2.3.1). A cross section of the mid colon, which contained part of a fecal pellet to better visualize the mucus layer, was also taken for alcian blue/periodic acid-Schiff (PAS) staining (as described in section 2.3.3).

### H&E and alcian blue/PAS staining

The longitudinal distal colonic sections, used for H&E staining, were fixed in 10% neutral buffered formalin (Fischer Scientific) for 24 hours and then transferred to 70% ethanol. The cross sections of the mid colon, used for alcian blue/PAS staining, were immediately fixed in methacarn (60% methanol, 10% acetic acid glacial and 30% chloroform) at 4°C overnight, transferred to 100% methanol for 1 hour and then 100% ethanol. Fixed tissue was embedded in paraffin and 4 μm sections were cut.

### H&E staining

H&E staining was preformed using standard procedures. In brief, the slides were first washed in H_2_O for 1 minute and then stained with hematoxylin for 5 minutes. The slides were washed again in H_2_O for 1 minute and then dipped 1-2 times in 1% acid alcohol. After washing for 30 seconds the slides were added to saturated lithium carbonate for 30 seconds. The slides were washed again and then checked microscopically to ensure the nuclear staining was adequate. If necessary, the slides were re-stained by repeating the previous steps. The slides were next immersed in 70% isopropyl alcohol for 30 seconds and then stained with 1% eosin (in 80% alcohol) for 15-30 seconds. The slides were added to 80% and then 90% isopropyl alcohol for 10 seconds each, and then washed in 100% isopropyl alcohol twice for 1 minute each. They were allowed to air dry and then a cover slip was added using mounting media.

### Histopathological scoring

H&E stained sections were blinded and imaged at 20x magnification. Two blinded scorers used a modified version of a previously described histopathological scoring system^27^ to assess disease severity using six histopathological criteria: 1) inflammatory cell infiltrate, 2) goblet cell loss, 3) crypt density loss, 4) crypt hyperplasia, 5) external muscle layer thickening and 6) submucosal infiltration. The average of the scores of the two blinded scorers is reported.

### Alcian blue/PAS staining

Slides were deparaffinized and rehydrated in distilled H_2_O. The tissue sections were stained in alcian blue 8GX (pH 2.5) for 30 minutes and the slides were then washed in tap H_2_O for 5 minutes. The slides were placed in periodic acid solution for 10 minutes and washed in tap water for an additional 5 minutes. The slides were then placed in Coleman’s Schiff reagent solution for 10 minutes and washed in lukewarm distilled H_2_O for 10 minutes. Lastly, the tissue sections were dehydrated using 95% ethanol, 100% ethanol and xylene – each twice for 2 minutes, and the slides were mounted using resinous medium.

### Mucus layer thickness and goblet cell counting

Alcian blue/PAS stained sections were blinded and imaged at 40x magnification. Between 7 and 10 images of each section of mid colonic tissue were taken. The mucus layer thickness and goblet cell numbers were evaluated using ImageJ (version 1.52q). For each image the average of 3 mucus layer thickness measurements and the average counts of goblet cells from 5 full length, well oriented crypts were obtained.

### T cell subset analysis

Spleens and mesenteric/colonic lymph nodes (MLN) were harvested from euthanized mice, processed into single cell suspensions and stimulated with 50 ng/mL of phorbol 12-myristate 13-acetate (PMA, Sigma) and 1 μg of ionomycin (Sigma) for 4h in the presence of 1ul/mL of Golgi-Plug (BD Biosciences). After stimulation, cells were Fc-blocked with anti-CD16/32 antibody (93, Biolegend), viability stained (eFluor780, Thermofisher) and surfaced stained for anti-Thy1.1 (HIS51, Thermofisher), CD4 (GK1.5, Thermofisher), and CD25 (PC61, Thermofisher). Following surface staining, cells were fixed with Fixation/Permeabilization buffer (Thermofisher) and intracellularly stained for the presence of T-bet (4B10, Thermofisher), Rorγt (AFKJS-9, Thermofisher), IL-10 (JES5-16E3, Thermofisher) IL-17A (eBio17B7, Thermofisher), IL-22 (IL-22JOP, Thermofisher), IFNγ (XMG1.2, Thermofisher), and TNF (MP6-XT22, Thermofisher). Samples were analyzed on a 4-laser LSR II flow cytometer (BD Biosciences) and FlowJo software, version 10.7.1 (Tree Star, Ashland, OR).

### Gut microbiome analysis

Stool samples were collected from mice the day of adoptive T cell transfer (pre-intervention) as well as the day prior to the end of experiment (post-intervention). Cecal contents were collected at the end of the experiment for short-chain fatty acid (SCFA) analysis. The stool and cecal samples were stored at −80°C until analysis.

### DNA extraction and quantification

Bacterial genomic DNA was extracted from stool following the Omega E.N.Z.A^®^ Soil DNA kit instructions with a few minor modifications. In brief, 65 mg ± 5% of stool was weighed out into bead beating tubes. The stool was homogenized at 30 m/s for 3 minutes with bead beating solution. The DNA was eluted in 50 μl of elution buffer while DNA concentrations were quantified using both a NanoDrop One (Thermofisher) spectrophotometer and Qubit™ 4 fluorometer (Thermofisher).

### Gut microbiota analysis using droplet digital PCR

The extracted bacterial DNA was diluted 5000x, 500x, 200x and 100x prior to quantifying total bacteria, as well as *Bifidobacterium* spp., *Bacteroides* spp. and *Faecalibacterium prausnitzii;* respectively. The primers and specific annealing temperatures used are summarized in Table 2. The Biorad QX200 droplet digital PCR (ddPCR) system was used to generate absolute quantification data. PCR was performed using 2 μL of template DNA and the following reagents: 11 μL of Evagreen Supermix, 0.5 μL of forward primer (10 μM), 0.5 μL of reverse primer (10 μM) and 11 μL of PCR grade H_2_O. Prior to undertaking PCR, the template DNA and reagents were portioned into up to 10,000 droplets using an automated droplet generator. Once droplets were formed, the PCR amplification was performed using the Biorad C1000 Touch Thermal Cycler under the following conditions: Enzyme activation at 95°C for 5 minutes, 40 cycles of denaturation (95°C for 3 seconds), annealing/extension (refer to primer specific temperatures in Table 2); for 1 minute and finished with signal stabilization (90°C for 5 minutes). The ramp rate used was 2.5°C per second. The droplet reader and associated QuantaSoft software were used to quantify the concentrations of total bacteria, *Bifidobacterium* spp., *Bacteroides* spp. and *F. prausnitzii* present in each sample.

**Table 2:**
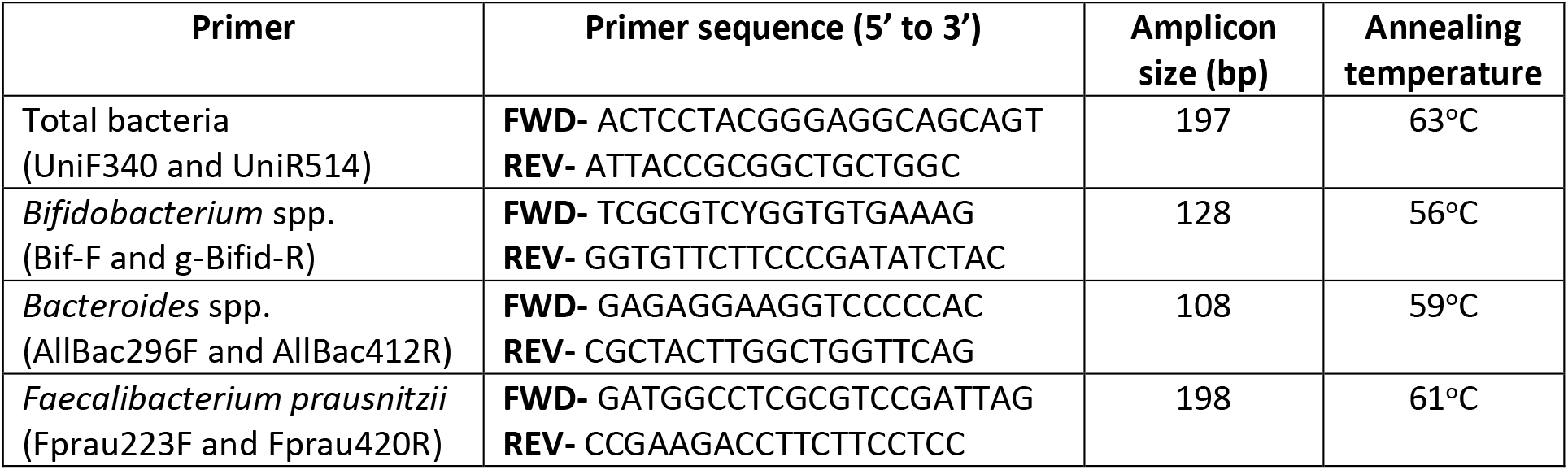
Forward and reverse primer sequences, amplicon size and annealing temperatures of the ddPCR primers.

### Short-chain fatty acid analysis using gas chromatography

The SCFA extraction method used is identical to the procedure published by Zhao and colleagues^28^. In brief, cecal samples were resuspended in MilliQ-grade H_2_O, and homogenized using MP Bio FastPrep for 1 minute at 4.0 m/s. 5M HCl was added to cecal suspensions to a final pH of 2.0. Acidified cecal suspensions were incubated and centrifuged at 10,000 rpm to separate the supernatant. Cecal supernatants were spiked with 2-Ethylbutyric acid for a final concentration of 1 mM. Extracted SCFA supernatants were stored in 2 ml GC vials, with glass inserts. SCFA were detected using gas chromatography (Thermo Trace 1310), coupled to a flame ionization detector (Thermofisher). The SCFA column used was the Thermofisher TG-WAXMS A GC Column (30 m, 0.32 mm, 0.25 um). The following settings were used for detection – Flame ionization detector: temperature – 240°C, hydrogen – 35 mL/min, Air – 350 mL/min, makeup gas (nitrogen) – 40 mL/min; Inlet: carrier pressure – 225 kPa, column flow – 6 mL/min, purge flow – 5 mL/min, split flow – 12 mL/min, splitless time – 0.75 min; Oven: temperature gradient – 100-180°C, gradient time – 10 min.

### Statistical analysis

One-way ANOVA was used to determine whether there were any significant differences in T cell subsets (only the chow, EEN and EEN-IN groups), spleen weight, colon length, body weight, disease activity score, SCFAs, goblet cell numbers and mucus thickness between the 4 intervention groups. Two-way ANOVA was used to determine if there were any significant differences in histopathology scores and pre- and post-intervention gut microbiota between the 4 intervention groups. GraphPad Prism software, version 8 for macOS, was used to perform all statistical analyses. All results presented are expressed as the mean values ± the standard errors (SEM). A p-value of 0.05 or less was considered significant with asterisks indicating significant differences in the figures. All authors had access to the study data and have reviewed and approved the final manuscript.

## Conflict of interest

All authors declare that they have no conflicts of interest.

## Funding statement

This work was supported by discovery grants (2016-05338 [KJ], 2018-05120 [BAV]) from the Natural Sciences and Engineering Research Council, the Children with Intestinal and Liver Disorders (CHILD) Foundation (KJ), operating grants (PJT-148846 and 159528 [BAV]) from the Canadian Institutes of Health Research (CIHR) and Crohn’s and Colitis Canada, and a Trainee Award from the Michael Smith Foundation for Health Research (17829 [GRH]).

## Abbreviations

BCFA: branch-chain fatty acids
CD: Crohn’s disease
DAS: disease activity score
ddPCR: droplet digital PCR
EEN: exclusive enteral nutrition
EEN-IN: inulin-type fructan enriched exclusive enteral nutrition
GI: gastrointestinal tract
GPR: G-protein coupled receptor
HDAC: histone deacetylase
IBD: inflammatory bowel disease
IN: inulin-type fructan
IP: intraperitoneal
MLN: mesenteric lymph nodes
NF-κB: nuclear-factor-kappa B
PAS: periodic acid-Schiff
PBS: phosphate buffered saline
PPARy: proliferator-activated receptor gamma
SCFA: short-chain fatty acid
Tregs: regulatory T cells.

## Notes

### Competing Interest Statement

The authors have declared no competing interest.

